# Deep learning-based imaging classification identified cingulate island sign in dementia with Lewy bodies

**DOI:** 10.1101/592865

**Authors:** Tomomichi Iizuka, Makoto Fukasawa, Masashi Kameyama

## Abstract

The differentiation of dementia with Lewy bodies (DLB) from Alzheimer’s disease (AD) using brain perfusion single photon emission tomography is important but has been a challenge because these conditions have common features. The cingulate island sign (CIS) is the most recently identified specific feature of DLB for a differential diagnosis. The present study aimed to examine the usefulness of deep learning-based imaging classification for the diagnoses of DLB and AD. We also investigated whether CIS was focused by the deep convolutional neural network (CNN) during differentiation.

Brain perfusion single photon emission tomography images were acquired from 80 patients each with DLB and with AD and 80 individuals with normal cognition (NL). The CNN was trained on brain surface perfusion images. Gradient-weighted class activation mapping (Grad-CAM) was applied to the CNN for visualization of the features that the trained CNN focused on.

Binary classifications between DLB and NL, DLB and AD and AD and NL were 94.69%, 87.81% and 94.38% accurate, respectively. The CIS ratios closely correlated with softmax output scores for DLB-AD discrimination (DLB/AD scores). The Grad-CAM highlighted CIS in the DLB discrimination. Visualization of learning process by guided Grad-CAM revealed that CIS became more focused by the CNN as the training progressed. DLB/AD score was significantly associated with three core-features of DLB.

Deep learning-based imaging classification was useful not only for objective and accurate differentiation of DLB from AD but also for predicting clinical features of DLB. The CIS was identified as a specific feature during DLB classification. The visualization of specific features and learning process could have important implications for the potential of deep learning to discover new imaging features.

## Introduction

Neuroimaging has contributed to the classification of neurodegenerative dementias such as dementia with Lewy bodies (DLB) and Alzheimers disease (AD). Early diagnoses of DLB and AD are important from prognostic and therapeutic perspectives and to distinguish between them is clinically vital. Disease-specific features have been extracted from brain perfusion single photon emission tomography (SPECT) images to assist with differential diagnoses of DLB and AD. Brain surface perfusion images produced by three-dimensional stereotactic surface projection (3D-SSP) [1] have been widely applied to statistical analysis that supported diagnoses of DLB and AD. A perfusion decrease in the parietal association cortex (PAC) and perfusion preservation in primary motor and primary somatosensory cortex are common in patients with DLB and AD [2, 3], which has interfered with distinguishing DLB from AD on perfusion SPECT images. An imaging feature for DLB discrimination is occipital hypoperfusion [4, 5, 6, 7]. Another finding that can produce difference between DLB and AD is perfusion in the posterior cingulate cortex (PCC). Hypoperfusion in PCC is observed in the early stage of AD, whereas the PCC is relatively preserved in DLB. The phenomenon of sparing of the PCC relative to the precuneus plus cuneus, which was termed the cingulate island sign (CIS) [8] has recently received focus, because it reflects concomitant AD pathology that impacts the clinical symptoms of DLB[9, 10]. We found that the CIS peaks at the stage of mild dementia and gradually disappears as DLB progress [11]. Thus, the CIS can help to differentiate DLB from AD especially at the early stage [8, 12] with some exceptions including posterior cortical atrophy[13].

Recent advances in deep learning, a main branch of artificial intelligence that has a deep convolutional neural network (CNN) capable of automatic feature extraction from data, have remarkably improved the performance of image classification and detection [14, 15]. Some algorithms based on deep learning have been proposed to recognize or differentiate AD and mild cognitive impairment (MCI) [16, 17]. In contrast, the ability of a CNN to discriminate DLB has not been investigated in detail. Furthermore, a deep learning-based SPECT interpretation system that could differentiate DLB and AD has not been described. The biggest disadvantage of deep learning is that the imaging features used by the CNN for classification have remained unknown. However, gradient-weighted class activation mapping (Grad-CAM) can produce “visual explanations” from a CNN, which allows visualization of areas where a CNN focuses[18, 19].

The present study aimed to objectively and automatically classify brain surface perfusion images via 3D-SSP of DLB, AD and individuals with normal cognition (NL) using deep 2D-CNN. We also investigated whether a trained CNN can identify the CIS, which is the most recently recognized imaging feature of DLB. Furthermore, visualization of learning process was performed during training of the CNN.

## Results

### Deep CNN could accurately classify brain surface perfusion images

Table 1 summarizes the demographic and cognitive findings of 80 persons each with AD, DLB and NL. The deep CNN was applied to images (*n* = 160) including right-left flipped images from each group of 80 patients for binary classification (Figure 4). The accuracy of the classification was calculated by 10-fold cross validation. The binary differentiations between DLB and NL (DLB-NL) and DLB and AD (DLB-AD) and AD and NL (AD-NL) were 94.69±4.90%, 87.81±7.85% and 94.38±5.47% accurate (mean ± standard deviation), respectively. The AUCs of the ROC for differentiating DLB-NL, DLB-AD and AD-NL were 98.5%, 93.9% and 97.6% accurate, respectively.

**Table 1:** Demographic features of study participants

|  | DLB | NL | AD |
| --- | --- | --- | --- |
| Participants ($n$) | 80 | 80 | 80 |
| Age (y) | $77.7 \pm 6.3$ | $77.1 \pm 6.8$ | $78.0 \pm 4.9$ |
| Sex (M/F) | 44/36 | 40/40 | 36/44 |
| MMSE score | $22.8 \pm 1.3^*$ | $29.5 \pm 0.6$ | $22.4 \pm 1.9^*$ |
| SPECT images ($n$) | 160 | 160 | 160 |

Data are shown as numbers or means ± standard deviation. $*p < 0.05$ : Tukey-Kramer test compared with NL (two-sided). DLB, dementia with Lewy bodies; NL, normal cognition; AD, Alzheimer’s disease; MMSE, mini-mental state examination; SPECT, single photon emission computed tomography.

## The CIS ratios significantly correlated with DLB/AD and DLB/NL scores

Close and weakly significant correlations were found between CIS ratios and scores for DLB/AD (*r* = 0.546, *p <* 0.001; Figure 1a) and DLB/NL (*r* = 0.267, *p <* 0.05; Figure 1b) in patients with DLB. Thus, the CIS ratio contributed more to the differentiation of DLB-AD than of DLB-NL.

**Figure 1:**
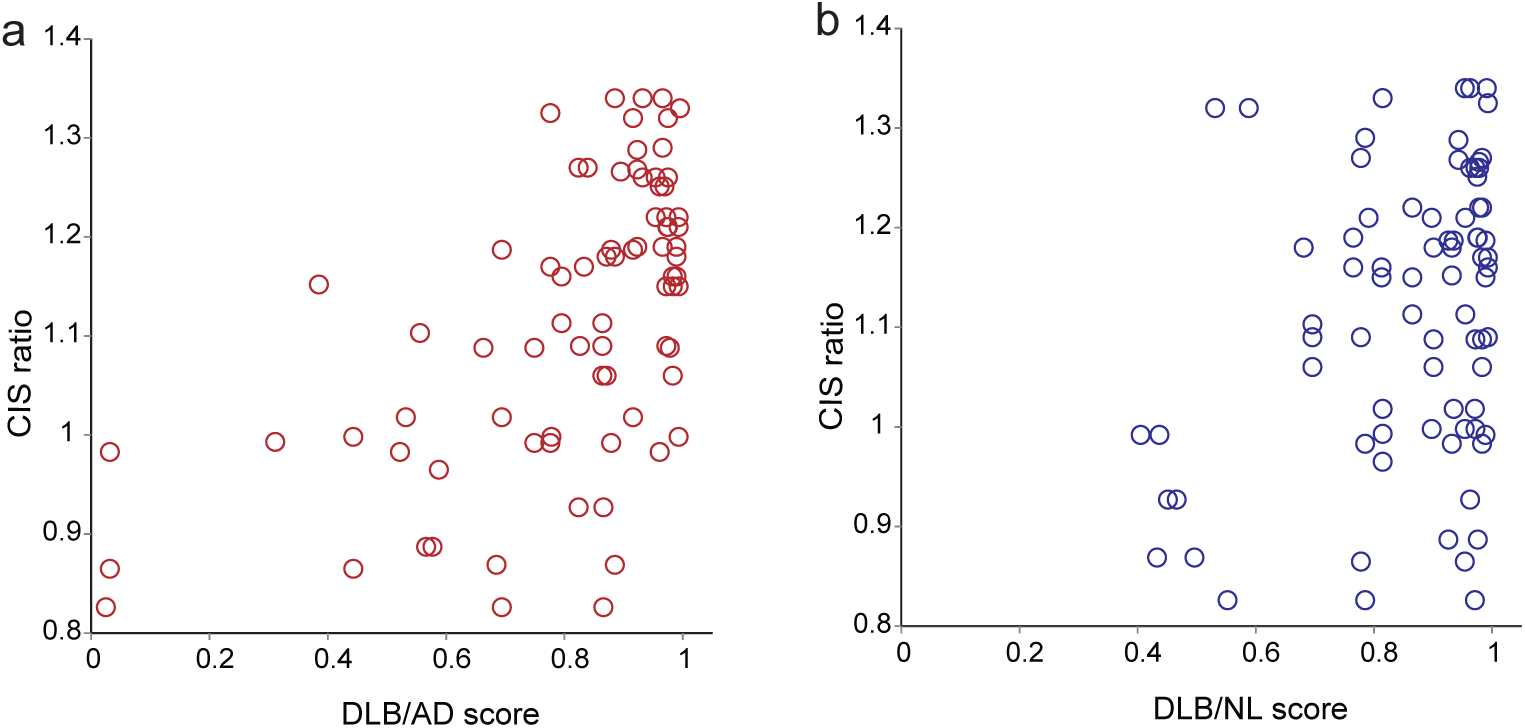
Association of CIS ratios with (a) DLB/AD and (b) DLB/NL scores. CIS ratio, DLB/AD score and DLB/NL score in patients with DLB were 1.11 0.14, 0.809 0.223 and 0.859 0.160, respectively (mean standard deviation). CIS ratios correlated closely with (a) DLB/AD scores (*r* = 0.546, *p <* 0.001) and weakly with (b) DLB/NL scores (*r* = 0.267, *p <* 0.05). CIS, cingulate island sign; DLB, dementia with Lewy bodies; AD, Alzheimer’s disease; NL, normal congition.

### Trained CNN identified CIS for DLB detection

Grad-CAM was applied to the trained CNN to produce heatmaps and guided Grad-CAM images for DLB-AD and DLB-NL discrimination. Heatmap clearly highlighted CIS in DLB to discriminate DLB and AD (Figure 2a). Guided Grad-CAM also had a limited range on image that focused on CIS. Brain perfusion images with obvious occipital hypoperfusion without CIS were also correctly labeled as DLB. In such images, Grad-CAM mostly highlighted cerebellum but not cortex. Heatmap and guided Grad-CAM for AD highlighted occipital lobe and cerebellum, but not PCC (Figure 2b). The CIS was highlighted less intensely in DLB-NL, than in DLB-AD discrimination (Figure 2c). Heatmap and guided Grad-CAM for NL diffusely highlighted occipital lobe, middle cingulate cortex, PCC and cerebellum (Figure 2d).

**Figure 2:**
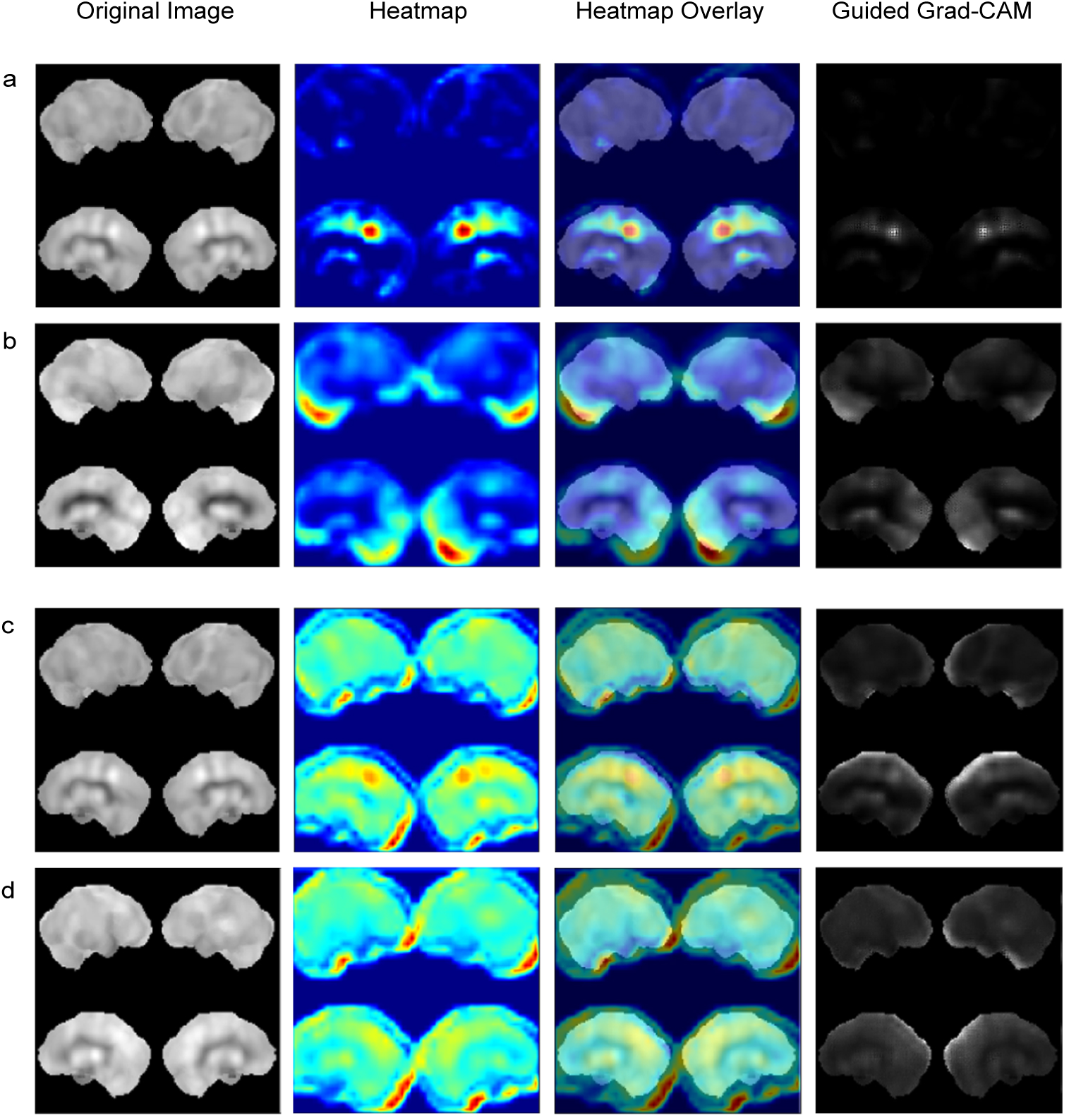
Visualization of features that the trained CNN recognized. Grad-CAM was applied to CNN trained with 100 epochs and produced heatmap, heatmap overlay and guided Grad-CAM. Original and Grad-CAM images from one patient with DLB in the DLB-AD (a) and DLB-NL (c) discrimination, respectively. Original and Grad-CAM images from a patient with AD in the DLB-AD discrimination (b). Original and Grad-CAM images from an individual with NL in the DLB-NL discrimination (d). Original images of (a), (b), (c) and (d) were correctly predicted. CNN, convolutional neural network; DLB, dementia with Lewy bodies; AD, Alzheimer’s disease; NL, normal congition; Grad-CAM, gradient-weighted class activation mapping.

### Visualization of feature extraction in the learning process of CNN

Grad-CAM visualized learning process to extract features that were useful for differentiation by showing altered images (Figure 3). In CNN trained for DLB-AD discrimination with 20 epochs, guided Grad-CAM and original images remained similar, indicating that CNN could not yet detect specific features. After 60 training epochs, guided Grad-CAM images became narrower and contrast became more obvious. After training with 100 epochs, CNN focused more tightly on CIS in DLB (Figure 3ab) and occipital lobe, cerebellum and sensorimotor area in AD (Figure 3cd).

**Figure 3:**
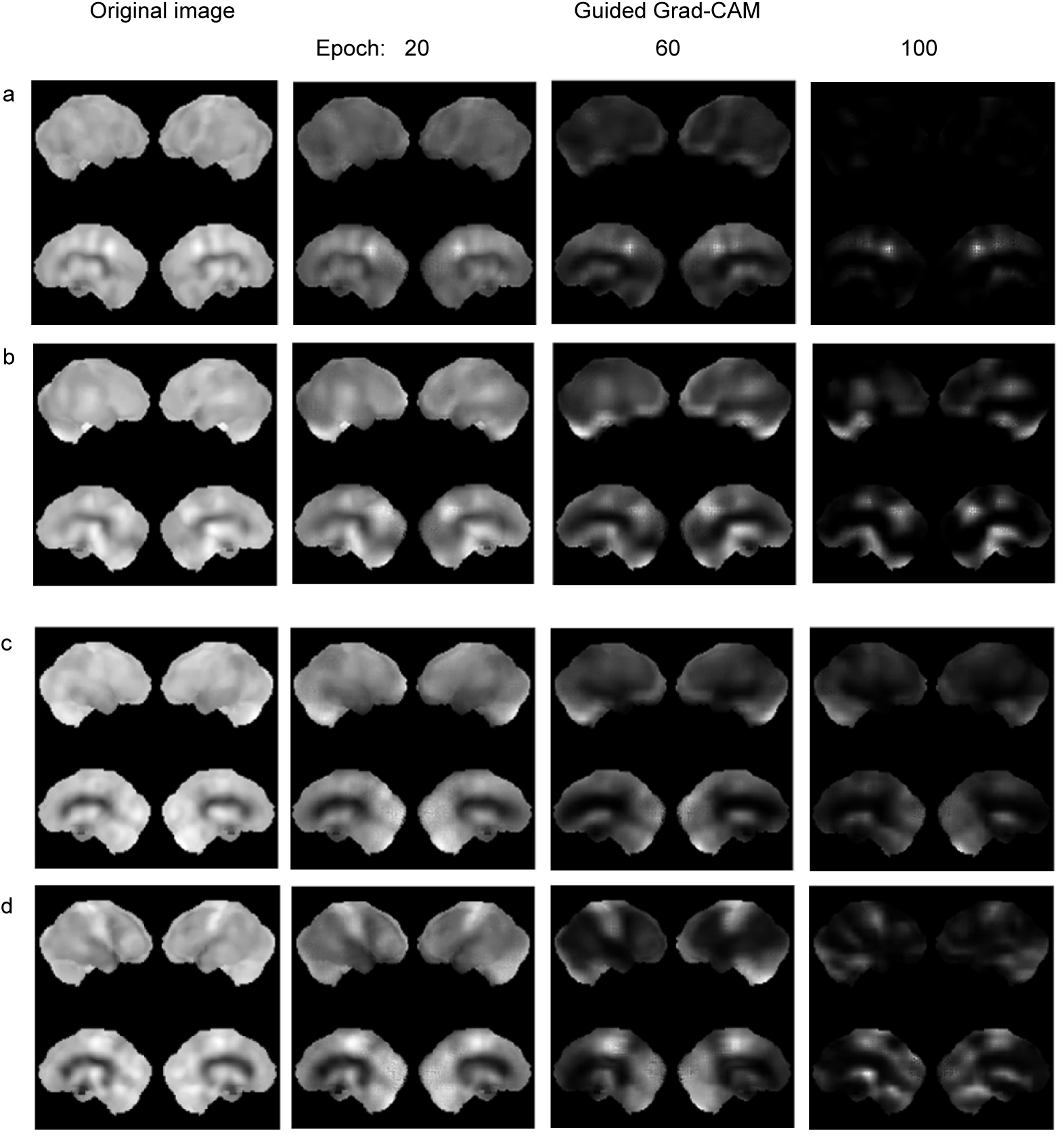
Alteration of guided Grad-CAM images in the learning process. Original and guided Grad-CAM images are from two patients each with DLB and AD. Two patients each with DLB (a) and (b), and AD (c) and (d). Training accuracies at 20, 60 and 100 epochs were 0.7682, 0.8922 and 0.9850, respectively. Validation accuracies at 20, 60 and 100 epochs were 0.6250, 0.7500 and 0.8750, respectively. Thus, 100 epochs was regarded as appropriate for training. The guided Grad-CAM images of both DLB and AD became reduced with increasing number of epochs. Original images of (a), (b), (c) and (d) were correctly predicted. CIS, cingulate island sign; DLB, dementia with Lewy bodies; AD, Alzheimer’s disease; Grad-CAM, gradient-weighted class activation mapping.

### DLB/AD score was associated with core-features of DLB

Association between neuroimaging indices (*i.e.*, CIS ratio, DLB/AD and DLB/NL score) and clinical symptoms (*i.e.*, four core-features and verbal memory) were analyzed. DLB/AD score was significantly correlated with hallucination, parkinsonism and RBD, but not with fluctuation (Table 2). In contrast, DLB/NL score was not correlated with any of them. The CIS ratio was correlated with hallucination and RBD. DLB/AD score and CIS ratio were also significantly correlated with verbal memory.

**Table 2:** Association between neuroimaging indices and clinical symptoms of DLB

|  | Correlation coefficient |  |  |
| --- | --- | --- | --- |
|  | CIS ratio | DLB/AD score | DLB/NL score |
| Hallucination | 0.307* | 0.235* | 0.203 |
| Fluctuation | 0.148 | 0.117 | 0.078 |
| Parkinsonism | 0.104 | 0.319* | 0.212 |
| RBD | 0.450** | 0.268* | 0.091 |
| Verbal memory | 0.611** | 0.487** | 0.201 |

\* $p < 0.05$ ; \*\* $p < 0.01$ : Spearman rank correlation coefficients (two-tailed). CIS, cingulate island sign; DLB, dementia with Lewy bodies; AD, Alzheimer’s disease; NL, normal cognition; RBD, REM sleep behavioral disorder.

## Discussion

Our CNN identified the CIS as an imaging feature during DLB-AD discrimination. The CIS ratios closely correlated with DLB/AD scores. Furthermore, heatmaps generated by the Grad-CAM highlighted the CIS in DLB. The guided Grad-CAM also focused on the CIS and became restricted to the CIS as the learning process progressed. The indirect evidence of the correlation coefficients might show that typical DLB has a higher CIS ratio. However, the trained CNN automatically and objectively identified the CIS as an important feature of DLB prediction, considering that the Grad-CAM could visualize the target of CNN for the classification. The present findings defined the potential of deep learning to discover new features in the field of image diagnosis.

Deep CNN could accurately classify brain surface perfusion images. The classification accuracies of DLB-NL, DLB-AD and AD-NL were 94.69%, 87.81% and 94.38%, respectively. Most previous studies using deep learning-based classification aimed to diagnose AD and MCI but not DLB using 3D-CNN, and the CNN diagnosis of DLB using FDG PET or perfusion SPECT has never been reported. Suk *et al.*[17] showed that the mean accuracies of MRI, FDG PET and MRI+PET with 3D-CNN were 92.38%, 92.20% and 95.35%, respectively. Liu *et al.*[16] generated accuracies of 90.18% (MRI), 89.13% (PET) and 90.27% (MRI+PET). Our 2D-CNN with brain surface perfusion images extracted from whole brain perfusion SPECT data reached comparable discriminative accuracy. The distribution on brain perfusion and glucose metabolism images was similar [20]. Brain surface perfusion images represent extracted features that are useful for discriminating neurodegenerative dementia. It provides us a bird’s-eye view. Furthermore, 3D-CNN needs much more calculation to converge more parameters than 2D-CNN. Thus, 2D-CNN with brain surface perfusion images classified more efficiently than 3D-CNN with whole brain images; our method, which can be operated in a standard computer has potential to prevail in clinical settings.

The CIS was more involved in the discrimination of DLB-AD rather than of DLB-NL, considering the higher correlation coefficients of the CIS ratios and DLB/AD scores than the CIS ratios and DLB/NL scores. The Grad-CAM supported this notion by focusing on the CIS as an imaging feature of DLB in the DLB-AD and DLB-NL discrimination. Heatmap and guided Grad-CAM highlighted the CIS in the DLB-AD discrimination, while CIS was less highlighted in the DLB-NL discrimination. As DLB and AD have common features such as rCBF decreases in the PAC, classification is usually more difficult for DLB-AD than DLB-NL. Most patients with DLB have concomitant AD pathology[21], which reportedly affects the CIS of patients with DLB. Specifically, the CIS is not obvious in DLB with abundant AD pathology. Similar to the CIS ratios, DLB/AD scores in DLB also reflect the degree of imaging features of AD that are presumably produced by concomitant AD pathology. Therefore, low CIS ratios and DLB/AD scores represent a high degree of concomitant AD pathology. Conversely, high CIS ratios and DLB/AD scores represent “pure” DLB. This explains why the CIS ratios had a good correlation with DLB/AD scores.

The Grad-CAM revealed that the CNN classified SPECT images in a manner unlike that of humans. Nuclear medicine physicians simultaneously evaluate these hypoperfused areas and preserved regions to differentiate DLB from AD and often consider contrast of preserved and decreased areas. In contrast, heatmaps generated by the Grad-CAM were placed only on regions with preserved rCBF in both AD and DLB in the appropriately trained CNN. Guided Grad-CAM images became narrower and restricted to more preserved regions as learning progresses. In line with these findings, the CNN focused only on preserved regions to classify brain surface perfusion images of both DLB and AD. Regardless of the manner of classification, the CNN still identified the CIS as an important imaging feature of DLB.

DLB/AD score was significantly correlated with scores of three core-features, namely hallucination, parkinsonism and RBD. In contrast, DLB/NL score was not correlated with any of them. The finding suggested that DLB/AD scores closely represented various symptoms of DLB. Similar to DLB/AD score, CIS ratio was also correlated with hallucination and RBD. As CIS is reportedly reflects AD pathology, close correlation of CIS ratio with DLB/AD score indicated that DLB/AD score also reflects comorbid AD pathology. Hallucination was frequently observed in DLB without AD pathology [22]. Manifestation of RBD was reportedly associated with less severe concomitant AD pathology[23]. Our finding was consistent with the previous reports showing the association between core-features and AD pathology. Furthermore, DLB/AD score was also correlated with verbal memory score, which reflects the fact that memory impairment is prominent in patients with AD rather than those with DLB. Thus, DLB/AD score was useful not only for the discrimination but also for predicting clinical features of DLB.

Our deep learning system would be beneficial to health care finance. Dopamine transporter (DaT) imaging[24] and [^123^I] MIBG cardiac sympathetic nerve scintigraphy[25] are authentic in clinically discriminating DLB from AD and the DLB guidelines treat DaT imaging and [^123^I] MIBG scintigraphy as indicative biomarkers[26]. However, to assess all amnestic pa-tients using two more nuclear medicine examinations might be too costly. Brain perfusion SPECT is more commonly used to detect AD, especially when a diagnosis is uncertain. Consequently, we suggest that our diagnostic system and perfusion SPECT could be initially applied to investigate DLB in patients with suspected AD before using DaT and cardiac sympathetic nerve imaging.

This study has several limitations. Each group comprised only 160 augmented images from 80 individuals because this study proceeded at a single institution. However, our brain surface perfusion images were normalized by 3D-SSP and applied only to binary classification. Therefore, we considered that the accuracy was sufficient regardless of the limited number of patients. The accuracy of FDG PET might be better, but perfusion SPECT is more accessible, and it has been proven as a valid alternative in the absence of FDG PET [27] and images with [^123^I] IMP shows good contrast due to its high first-pass extraction[11, 28]. Recent CNN studies have attempted to enhance accu-racy using various combinations of imaging modalities [16, 17]. Although the ability of a 2D-CNN with brain surface perfusion images was comparable to previous findings with such combinations, future studies should examine combinations of perfusion SPECT with other imaging modalities to enhance accuracy.

## Methods

### Participants

Brain perfusion SPECT images of 80 persons each with DLB, AD and NL were included for diagnostic classification and CNN learning. Cognitive function was evaluated using the Clinical Dementia Rating and the Mini-Metal Status Examination (MMSE). Probable DLB and probable AD were diagnosed according to the McKeith criteria[26] and the criteria of the National Institute for Neurological and Communicative Diseases Alzheimer’s Disease and Related Disorders Association [29], respectively. Hallucination, fluctuation of consciousness, parkinsonism and REM sleep behavioral disorder (RBD) were assessed by Neuropsychiatric Inventory (NPI), Clinician Assessment of Fluctuation[30], United Parkinson’s Disease Rating Scale-Motor Score (UPDRS-MS) and Japanese version of the REM sleep behavior disorder screening questionnaire (RBDSQ-J)[31], respectively. Verbal memory was evaluated using sum of the five recall trials (1-5) of Ray Auditory Verbal Learning Test (RAVLT).

All procedures were approved by the Ethical Re-view Board at Fukujuji Hospital. We followed the clinical study guidelines of Fukujuji Hospital, which conformed to the Declaration of Helsinki (2013). We provided the healthy volunteers, patients and their families with detailed information about the study, and all provided written informed consent to participate.

### Brain perfusion SPECT imaging

Persons resting with their eyes closed and ears unplugged were assessed by SPECT using a Symbia Evo Excel with [^123^I] IMP, a gamma camera (Siemens Medical Solutions, Malvern, PA, USA) and fan beam collimators. Fifteen minutes after an intravenous infusion of [^123^I] IMP (167 MBq), SPECT images were acquired in a 128 × 128 matrix with a slice thickness of 1.95 mm (1 pixel) over a period of 30-40 min. The images were reconstructed by filtered back projection using a Butterworth filter, attenuation was corrected using the Chang method (attenuation coefficient = 0.1 cm^*-*1^) and scatter was corrected using a triple energy window. Brain surface perfusion images produced using 3D-SSP[1] were augmented by flipping from left to right. The regional cerebral blood flow (rCBF) in regions of interest (ROI) on the PCC, precuneus and cuneus was measured as described[11]. The mean value in the bilateral PCC ROI was divided by the mean value in the bilateral precuneus plus cuneus ROI to derive CIS ratios from [^123^I] IMP SPECT images.

### Preparation for deep convolutional neural network

Figure 4 summarizes the architecture of our deep CNN. The network was built with TensorFlow (Google, Mountain View, CA, USA), a deep learning framework. We did not use transfer learning to visualize the process of learning. After the convolution operation, rectified linear unit (ReLU) and max-pooling operations proceeded on the output of convolution. The ReLU kept positive input values whereas negative input values were changed to zeros. The max-pooling operation selected the maximum value and input this value into a smaller feature map. Input data were extracted from brain perfusion SPECT images. The input image had a matrix of 200 × 200 pixels, which is a composite of 2 lateral and 2 medial surface images. Input values of voxels were rescaled within a range of 0 to 255, and then the mean scalar value of each SPECT volume was subtracted. The images were passed through the first convolutional layer which produced 193 × 193 × 32 output images after the 8 × 8 × 32 convolutional filter.

**Figure 4:**
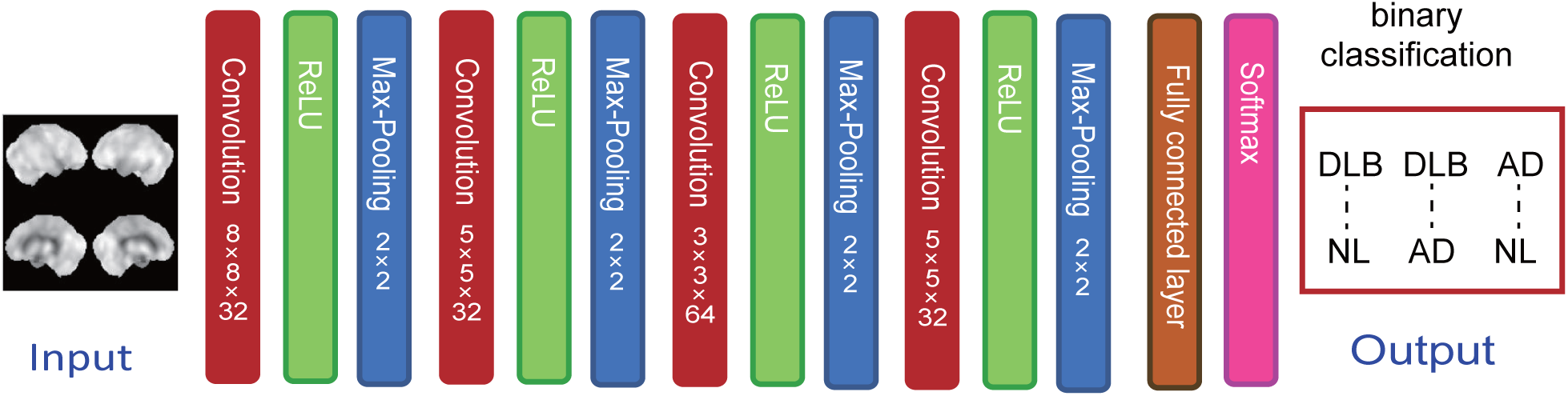
Architecture of deep convolutional neural network

Thereafter, ReLU activation and max pooling of a 2 × 2 pool proceeded. The second convolutional layer with a 5 5 32 filter and 92 92 32 output was followed by the ReLU activation and max-pooling layers. The third convolutional layer with a 3 3 64 filter and 44 × 44 × 64 output was followed by the ReLU activation and max-pooling layers. The last convolutional layer with a 5 × 5 × 32 filter and 18 × 18 × 32 output was followed by the ReLU activation and max pooling layers that produced a 9 9 32 output. Thereafter, a fully connected layer generated output, then a softmax function was applied to discriminate two labels. The softmax produces two numerical values of which the sum becomes 1.0. The output values for the binary differentiation of DLB-NL, DLB-AD and AD-NL are expressed as DLB/NL, DLB/AD and AD/NL scores, respectively. We employed binary discrimination to know if CNN recognizes CIS differently in discriminating DLB-AD and DLB-NL. The network was trained to minimize cross entropy losses between the predicted and true diagnoses based on the images. The CNN was trained for 100 epochs. The momentum parameter was and the learning rate was 0.0001. To visualize the decision made by the CNN, Grad-CAM was applied to the CNN. The Grad-CAM uses the gradients of any target flowing into the final convolutional network to produce heatmaps that highlight important regions upon which the CNN focuses. A guided Grad-CAM was created by fusing existing pixel-space gradient visualizations with the Grad-CAM to achieve both high-resolution and class-discrimination. We also used Grad-CAM to visualize learning process of the CNN trained with perfusion images.

## Statistics

The diagnostic and predictive accuracy of the CNN was calculated from 10.0% data from the training/validation set and 10-fold cross validation. An original images and its right-left flip image were in a same set of training or validation. Binary classification scores were evaluated using the receiver operating characteristic (ROC) curve analysis and area under the curve (AUC). Correlations between CIS ratios and DLB/AD or DLB/NL scores were assessed using Spearman rank correlation coefficients. All statistical analyses were performed with EZR (Saitama Medical Center, Jichi Medical University, Saitama, Japan), which is a graphical user interface for R (The R Foundation for Statistical Computing, Vienna, Austria). More precisely, it is a modified version of R commander designed to add statistical functions frequently used in biostatistics.

## Acknowledgements

This study was partly supported by Grant-in-Aid for Scientific Research (Kakenhi) Grant number 18K07488.

## Author contributions statement

T.I. contributed to conceptualization, patient recruitment, investigation, project administration and the initial draft manuscript preparation. T.I. and M.F. built CNN, analysed the data and produced figures. M.K. advised the project and revised the manuscript. All the authors discussed the project and have read and approved the final manuscripts.

## Additional information

### Competing financial interests

M.K. received research funding from Fujifilm RI Pharma, Nihon Med-Physics, and Daiichi-Sankyo.

